## Supporting Figures for "Rapid protein stability prediction using deep learning representations"

### Supporting Material for: Rapid protein stability prediction using deep learning representations

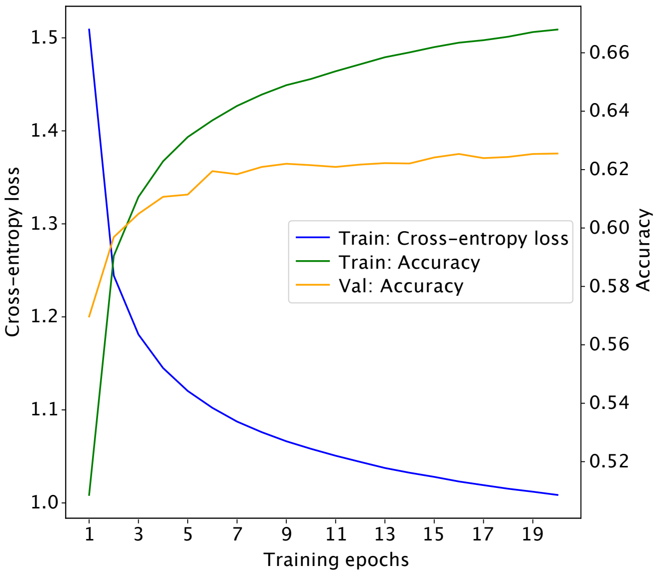


**Figure 2—figure supplement 1.** Learning curve for the self-supervised 3D convolutional neural network. The model obtained at epoch 15 achieves a classification accuracy of 63% on the validation set.

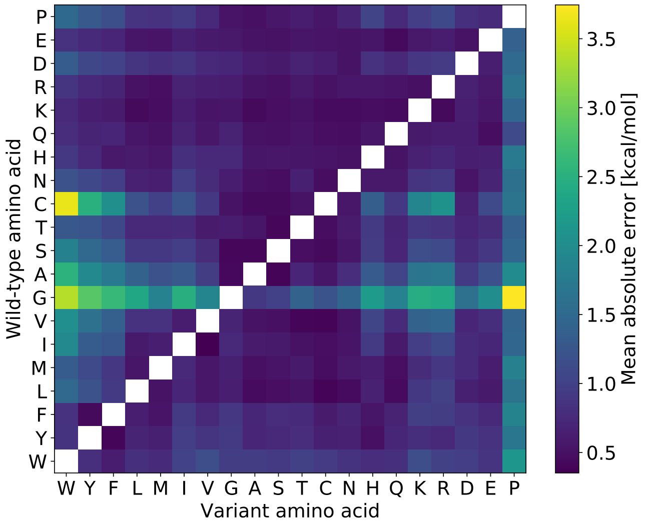


**Figure 2—figure supplement 2.** Mean absolute prediction error for RaSP on the validation set, split by amino acid type of the wild-type and variant residue. Substitutions from glycine and cysteine as well as to proline generally have higher errors.

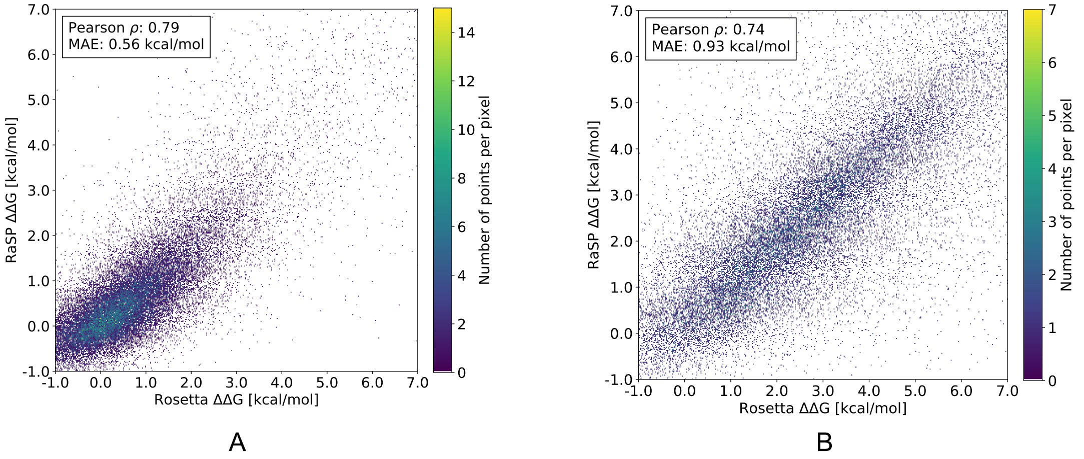


**Figure 2—figure supplement 3.** RaSP versus Rosetta $\Delta\Delta G$ values for a full saturation mutagenesis of 10 test proteins separated into either exposed (A) or buried (B) residues. We speculate, that the RaSP prediction task is harder in the case of buried residues because Rosetta $\Delta\Delta G$ values generally have higher variance in those regions. Pearson correlation coefficients and mean absolute errors (MAE) were for this figure computed using only variants with Rosetta $\Delta\Delta G$ values in the range [-1;7] kcal/mol. Buried and exposed residue were classified based a relative surface accessible surface area (SASA) cut-off of 0.2

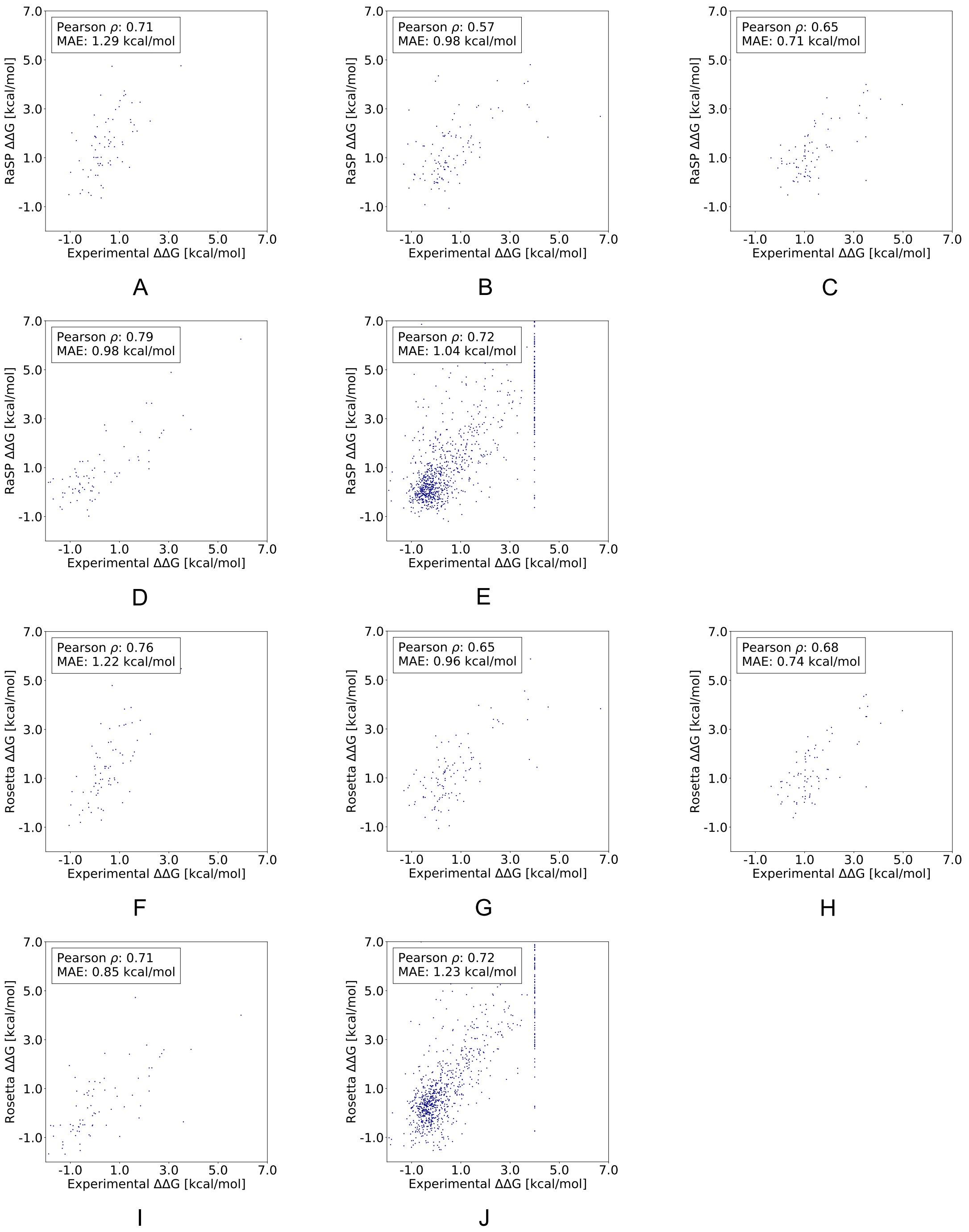


**Figure 3—figure supplement 1.** Comparing RaSP and Rosetta predictions to experimental stability measurements. Stability predictions obtained using (A–E) RaSP and (F–J) Rosetta are compared to experimental data for the five test proteins; myoglobin (1BVC), lysozyme (1LZ1), chymotrypsin inhibitor (2CI2), RNAse H (2RN2) and Protein G (1PGA)(***Kumar et al., 2006***; ***Ó Conchúir et al., 2015***; ***Nisthal et al., 2019***). In the experimental study of Protein G, 105 variants were assigned a $\Delta\Delta G$ value of at least 4 kcal/mol due to low stability, presence of a folding intermediate, or lack expression (***Nisthal et al., 2019***).

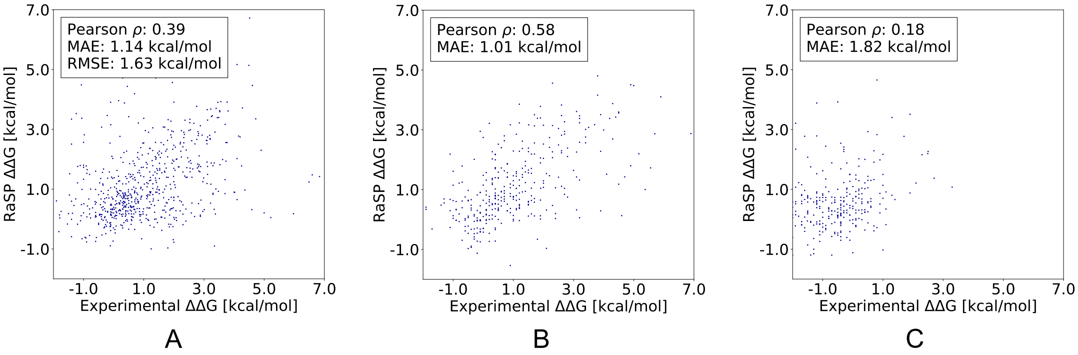


**Figure 3—figure supplement 2.** RaSP performance on three recently published data sets (***Pancotti et al., 2022***): (A) The S669 data set, (B) The Ssym+ direct data set, (C) The Ssym+ reverse data set.

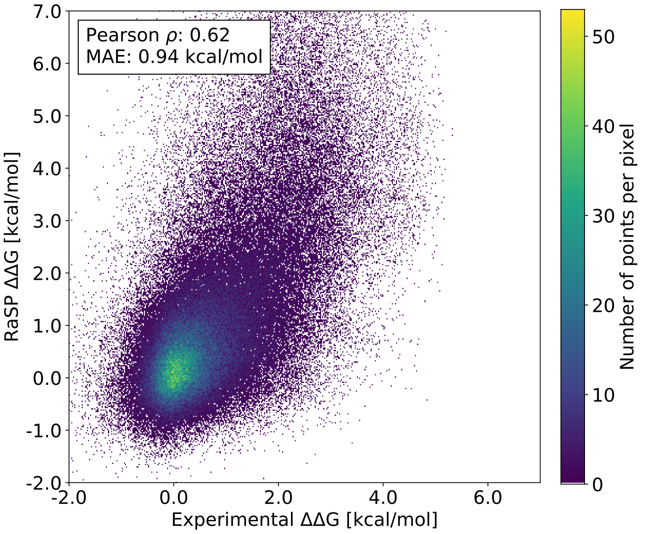


**Figure 3—figure supplement 3.** RaSP performance on the recently published mega-scale experiments (***Tsuboyama et al., 2022***). The experimental data has been filtered to include only well-defined experimental $\Delta\Delta G$ values from single substitution mutations in natural protein domains (***Tsuboyama et al., 2022***). This filtered data set contains a total of 164,524 variants across 164 protein domain structures.

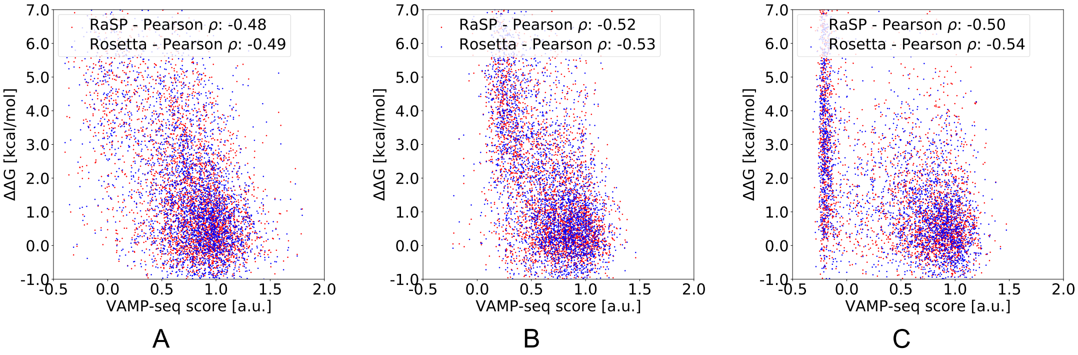


**Figure 3—figure supplement 4.** Benchmarking RaSP and Rosetta using VAMP-seq data. We compare stability predictions with VAMP-seq scores for three test proteins (A) TPMT (PDB: 2H11) (***Matreyek et al., 2018***), (B) PTEN (PDB: 1D5R) (***Matreyek et al., 2018***) and (C) NUDT15 (PDB: 5BON) (***Suiter et al., 2020***).

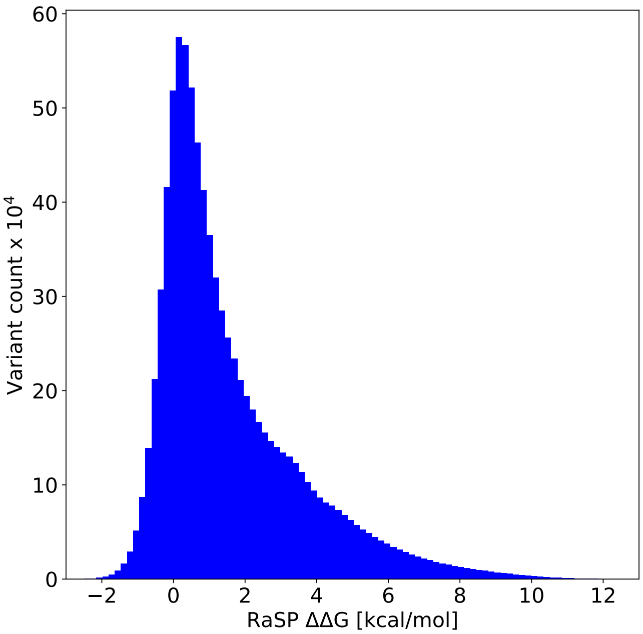


**Figure 5—figure supplement 1.** Histogram of $\Delta\Delta G$ values from saturation mutagenesis using RaSP on 1,366 PDB structures corresponding to ~8.8 million predicted $\Delta\Delta G$ values.

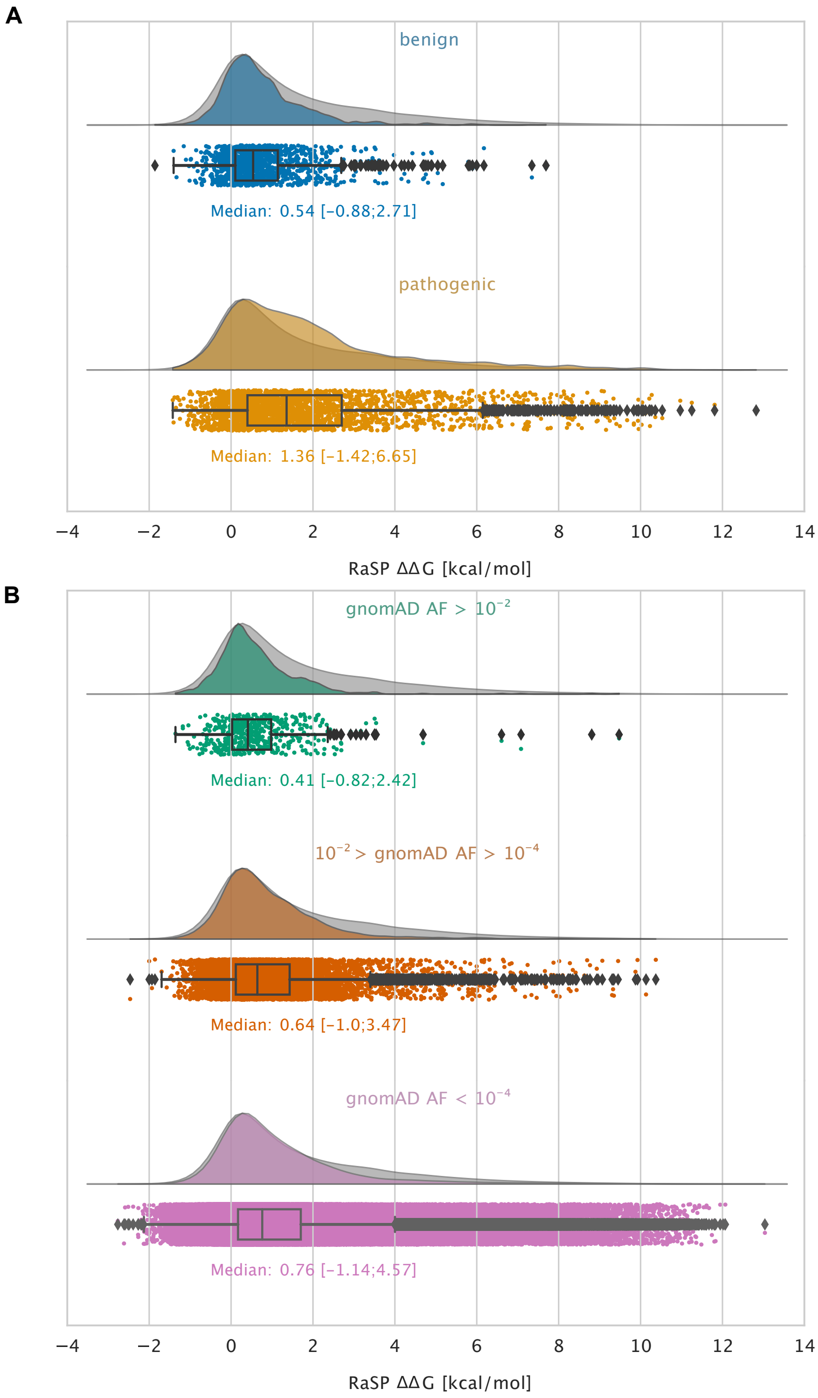


**Figure 5—figure supplement 2.** Large-scale analysis of disease-causing variants and variants observed in the population using the RaSP model. The grey distribution shown in the background of all plots represents the distribution of $\Delta\Delta G$ for all single amino acid changes in the 1,366 proteins that we analysed. Each plot is also labelled with the median $\Delta\Delta G$ of the subset analysed as well as a range of $\Delta\Delta G$ values that cover 95% of the data in that subset. (A) Distribution of RaSP $\Delta\Delta G$ values for benign (blue) and pathogenic (tan) variants extracted from the ClinVar database (***Landrum et al., 2018***). We observe that the median RaSP $\Delta\Delta G$ value is higher for pathogenic variants compared to benign variants. (B) Distribution of RaSP $\Delta\Delta G$ values for variants with different allele frequencies (AF) extracted from the gnomAD database (***Karczewski et al., 2020***) in the ranges i) AF > $10^{-2}$ (green), ii) $10^{-2}$ > AF > $10^{-4}$ (orange), iii) AF < $10^{-4}$ (purple). We observe a gradual shift in the median RaSP $\Delta\Delta G$ going from common variants (AF> $10^{-2}$) towards rarer ones (AF< $10^{-4}$).

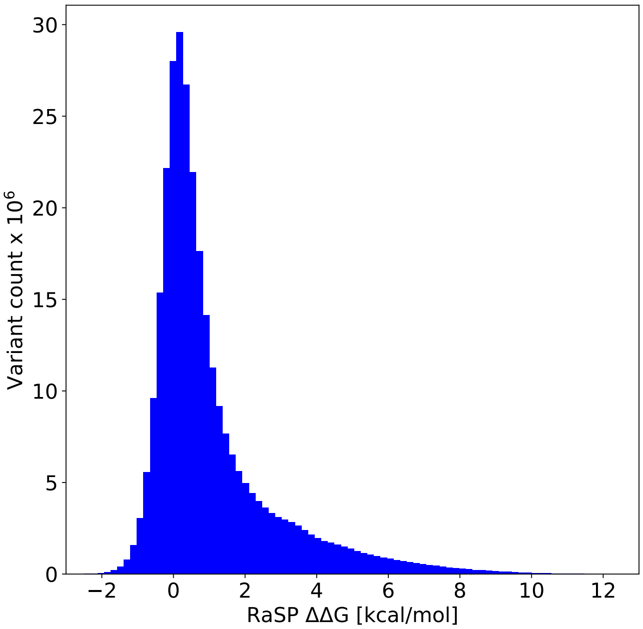


**Figure 5—figure supplement 3.** Histogram of $\Delta\Delta G$ values from saturation mutagenesis using RaSP on predicted structures of the entire human proteome corresponding to ∼300 million predicted $\Delta\Delta G$ values predicted from 23,391 protein structures.
